# The androgen receptor in mesenchymal progenitors regulates skeletal muscle mass via *Igf1* expression in male mice

**DOI:** 10.1101/2023.11.21.568190

**Authors:** Hiroshi Sakai, Hideaki Uno, Harumi Yamakawa, Kaori Tanaka, Aoi Ikedo, Akiyoshi Uezumi, Yasuyuki Ohkawa, Yuuki Imai

**Author notes:** Hiroshi Sakai, Yuuki Imai, Hiroshi Sakai, DVM, PhD, and Yuuki Imai, MD, PhD, Proteo-Science Center, Graduate School of Medicine, Ehime University, Shitsukawa, Toon, Ehime, 791-0295, Japan., (H.S.), (Y.I.). **Author Contributions:** Conceptualization, H.S. and Y.I.; Methodology, H.S., H.U., K.T., A.I., A.U., Y.O., and Y.I.; Software, H.S., H.U., and K.T.; Investigation, H.S., H.U., H.Y., K.T., and A.I.; Writing – Original Draft, H.S. and Y.I.; Writing – Review & Editing, H.S. and Y.I.; Funding Acquisition, H.S. and Y.I.; Resources, H.S., K.T., A.U., Y.O., and Y.I.; Supervision, H.S. and Y.I. **Competing Interest Statement:** The authors declare no competing interests.

## Abstract

Androgens exert their effects primarily by binding to the androgen receptor (AR), a ligand-dependent nuclear receptor. While androgens have anabolic effects on skeletal muscle, previous studies reported that AR functions in myofibers to regulate skeletal muscle quality, rather than skeletal muscle mass. Therefore, the anabolic effects of androgens are exerted via extra-myofiber cells or tissues. In this context, the cellular and molecular mechanisms of AR in mesenchymal progenitors, which play a crucial role in maintaining skeletal muscle homeostasis, remain largely unknown. In this study, we demonstrated expression of AR in mesenchymal progenitors and found that targeted AR ablation in mesenchymal progenitors reduced limb muscle mass in mature adult, but not young or aged, male mice, although fatty infiltration of muscle was not affected. The absence of AR in mesenchymal progenitors led to remarkable perineal muscle hypotrophy, regardless of age, due to abnormal regulation of transcripts associated with cell death and extracellular matrix organization. Additionally, we revealed that AR in mesenchymal progenitors regulates the expression of insulin-like growth factor 1 (Igf1), and that IGF1 administration prevents perineal muscles atrophy in a paracrine manner. These findings indicate that the anabolic effects of androgens regulate skeletal muscle mass via, at least in part, AR signaling in mesenchymal progenitors.

**Significance statement:** Androgens are essential not only for the development of male sexual characteristics but also for a range of physiological functions, including the regulation of skeletal muscle growth and function. Understanding the functionality of the androgen receptor (AR) is essential for comprehending the mechanisms through which androgens exert their effects on skeletal muscles, as these effects are mediated through AR binding. We demonstrate that AR is expressed in mesenchymal progenitors, which play a vital role in muscle homeostasis, and regulates the expression of insulin-like growth factor 1 (Igf1)—a key player in skeletal muscle growth—to control muscle mass. Our study provides significant insights into potential therapeutic strategies for addressing muscle atrophy conditions like sarcopenia.

## Introduction

Androgens are crucial regulators of various biological processes, including the development and maintenance of male characteristics. Notably, in a physiological context, the administration of androgens or androgen analogs has been shown to exert anabolic effects on muscles, resulting in increased muscle mass, strength, and body weight (1, 2). These physiological effects are mediated by activation of the androgen receptor (AR), a member of the nuclear receptor superfamily (3). However, despite that clinical administration of androgens in men can successfully increase skeletal muscle mass (4), the precise cellular and molecular mechanisms underlying the function of AR in muscle remain largely unexplored.

Skeletal muscle represents a complex tissue comprising multiple cell types, and the effects of androgens are likely mediated by various cell populations expressing AR within skeletal muscles. While male mice systemically lacking AR exhibit low skeletal muscle strength and mass, similar to female levels (5), we previously showed that muscle hypertrophy can be induced by androgen administration even in the absence of AR in myofibers, suggesting that non-myofiber AR contributes to the regulation of muscle mass (6). Furthermore, another of our studies demonstrated that deletion of AR in satellite cells, which are muscle stem cells responsible for muscle regeneration, had a limited effect on their proliferation and differentiation (7). Additionally, since limb muscle mass remains unaffected even in mice with AR ablation in both myofibers and muscle stem cells (8), other cells expressing AR within the muscles may play a role in regulating skeletal muscle mass.

Mesenchymal progenitors (9), also known as fibro/adipogenic progenitors (10), have been identified as a cell population distinct from myogenic cells characterized by the expression of platelet-derived growth factor receptor alpha (PDGFRα). These cells are involved in the maintenance and regeneration of skeletal muscle (11, 12). Conversely, mesenchymal progenitors have been implicated in pathological fibrosis and fat accumulation in skeletal muscle (13). AR expression has been reported in cells located in the interstitium or outside the basal lamina in both mouse and human skeletal muscles (14, 15). Additionally, ablation of AR specifically in embryonic cells of the mesenchyme led to impaired development of the perineal levator ani (LA) and bulbocavernosus (BC) muscles (16), which are particularly responsive to androgen. Furthermore, the reduced proliferation of undifferentiated myoblasts in these mutant mice suggested a potential paracrine regulatory role of mesenchymal AR in controlling skeletal muscles (16). Despite this knowledge, the specific role of AR in adult mesenchymal progenitors in skeletal muscles remains unexplored.

Here we investigate the role of AR expressed in mesenchymal progenitors in muscle maintenance using mice with mesenchymal progenitor-specific AR ablation. We show that AR in mesenchymal progenitors plays a critical role in maintaining skeletal muscle mass via regulation of insulin-like growth factor 1 (Igf1) expression in male mice.

## Results

### AR ablation in mesenchymal progenitors was validated in *PDGFRα-CreER;AR^L2/Y^* mice

AR expression in fibroblasts of the perineal LA muscles and in mesenchymal cells of the fetal LA muscles has been reported (14, 16, 17); however, it is unknown whether mesenchymal progenitors in adult limb muscle express AR. To examine the expression pattern of AR in mesenchymal progenitors, immunofluorescence staining of PDGFRα, a marker of mesenchymal progenitors (9), and AR was performed in adult (12 weeks old) male mice. We found that AR was expressed in mesenchymal progenitors staining positive for PDGFRα (Fig. 1A), as well as in myofibers surrounded by laminin-expressing basal lamina, consistent with previous findings (17, 18).

**Figure 1.**
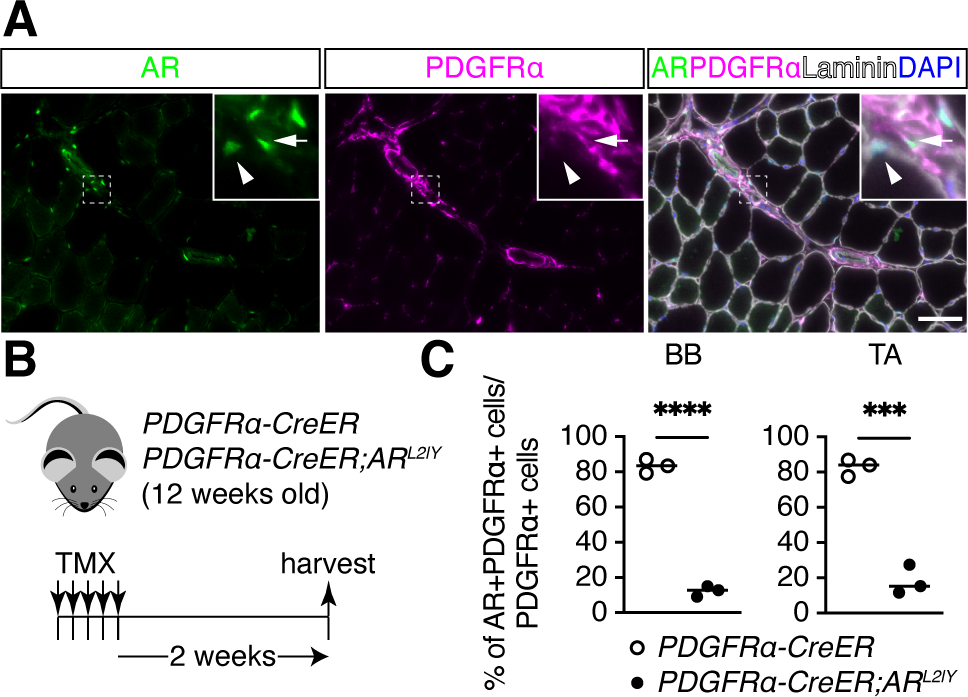
AR ablation in mesenchymal progenitors from *PDGFRα-CreER;AR^L2/Y^*mice. (A) Immunofluorescence staining of AR (left) and PDGFRα (middle) and of both together with laminin and DAPI staining (right) in male TA muscles. Arrows: AR+ and PDGFRα+ cells; arrowheads: AR+ nuclei surrounded by laminin. Scale bar, 50 µm. (B) Experimental design for tamoxifen (TMX) treatment followed by sample harvest. (C) Quantification of AR+ cells among PDGFRα+ cells in the biceps brachii (BB, left) and tibialis anterior (TA, right) muscles of control and mutant mice (n = 3). The two-sample test for equality of proportions: ***p < 0.001, ****p < 0.0001.

To investigate the role of AR in mesenchymal progenitors, *PDGFRα-CreER* (19) and *AR^L2/Y^* (20) mice were crossed to selectively delete the AR gene in mesenchymal progenitors expressing PDGFRα. Tamoxifen (TMX) was administered to Cre control male mice (*PDGFRα-CreER*) and mutant male mice (*PDGFRα-CreER;AR^L2/Y^*) for 5 consecutive days to induce AR ablation (Fig. 1B). To assess the efficiency of AR ablation, immunofluorescence staining of PDGFRα and AR was performed in control and mutant mice. Quantification of PDGFRα and AR expression in forelimb biceps brachii (BB) and hindlimb tibialis anterior (TA) muscles demonstrated a significant decrease of 90% in AR protein in mutant mice compared with control mice, confirming successful ablation of AR expression in mesenchymal progenitors from *PDGFRα-CreER;AR^L2/Y^* mice (Fig. 1C). In contrast, loss of AR in mesenchymal progenitors did not affect the number of PDGFRα+ cells (Fig. S1A). Taken together, these findings demonstrate successful and specific loss of AR expression in PDGFRα+ cells, rendering them an optimal model for investigating AR in mesenchymal progenitors in skeletal muscles.

### Lack of AR in mesenchymal progenitors had a limited effect on steady-state limb skeletal muscles

Because ablation of PDGFRα+ cells in intact muscles results in skeletal muscle atrophy (11, 12), we first analyzed the muscles of control and mutant mice under steady-state conditions at a young age (14 weeks old) (Fig. 1B). Body weight and limb skeletal muscle mass were not affected in mutant compared with control mice at 2 weeks after TMX administration (Fig. S1B). Moreover, the grip strength of mutant mice was comparable with that of control mice (Fig. S1C). The minimum Ferret diameter, which is the robust parameter in order to measure myofiber size, and number of myofibers in BB and TA muscles remained unchanged in mutant mice (Fig. S1D, E). As relayed signaling between mesenchymal progenitors and muscle stem (satellite) cells is critical for muscle hypertrophy (21), we also quantified the number of muscle stem cells.

Immunofluorescence staining of Pax7, a marker of muscle stem cells (22), revealed no difference in the number of muscle stem cells between control and mutant BB and TA muscles (Fig. S1F). We next investigated the fiber types of slow soleus muscles, which are affected by muscle fiber- and muscle stem cell-specific AR ablation (8, 23). Analysis of soleus muscles fiber types by immunofluorescence staining of type I (Myh7), type IIa (Myh2), and type IIx (without signals) muscle fibers demonstrated no difference in the fiber-type parameters (minimum Ferret diameter and frequency of each fiber-type) between control and mutant mice (Fig. S1G, H). Thus, AR expression loss in mesenchymal progenitors has a limited effect on the steady-state limb muscles of young mice.

### The impact of AR deficiency in mesenchymal progenitors on adipogenesis in muscles was limited

We also examined the adipogenesis of mesenchymal progenitors in mutant mice because adipocytes originate from mesenchymal progenitors in skeletal muscles (9). To address this, both the control and mutant mice were subjected to an 8-week regimen of high fat diet (HFD) feeding (Fig. S2A). No significant differences were observed between the control mice and mutant mice in terms of total body weight, hindlimb muscle weights, epididymal white adipose tissue (eWAT) mass, and subcutaneous white adipose tissue (sWAT) mass (Fig. S2B), and grip strength (Fig. S2C). Immunofluorescence staining of perilipin, a marker of adipocytes (24), along with laminin, demonstrated no significant difference in the minimum Ferret diameter of myofibers or the area occupied by adipocytes between the control and mutant TA muscles (Fig. S2D, E).

To further investigate the impact of AR deficiency in muscle adipogenesis, we employed a muscle injury model by administering glycerol injections into the TA muscle of both control and mutant mice (Fig. S2F). No significant differences were observed between the control and mutant mice in terms of the body and TA muscle weights at 2 weeks after injection (Fig. S2G). Immunofluorescence staining of perilipin in conjunction with laminin (Fig. S2H) revealed no significant difference in the minimum Ferret diameter of myofibers or the area occupied by adipocytes between the control and mutant TA muscles (Fig. S2I). These findings suggest that the absence of AR in mesenchymal progenitors does not significantly affect the adipogenic properties of TA muscles.

### AR deficiency in mesenchymal progenitors affected limb skeletal muscles in mature adult but not aged mice

To investigate the effect of the duration of AR ablation on mesenchymal progenitors, we analyzed mature adult (6 months old) mice and compared the Cre control group and mutant group (Fig. 2A). In the mature adult mutant mice, we observed decreases in body weight, TA, and gastrocnemius muscle weights (Fig. 2B), while there were no significant changes in the weights of eWAT and sWAT (Fig. 2B), which are affected by systemic AR ablation (25). However, grip strength (Fig. 2C) was comparable between the control and mutant mice.

**Figure 2.**
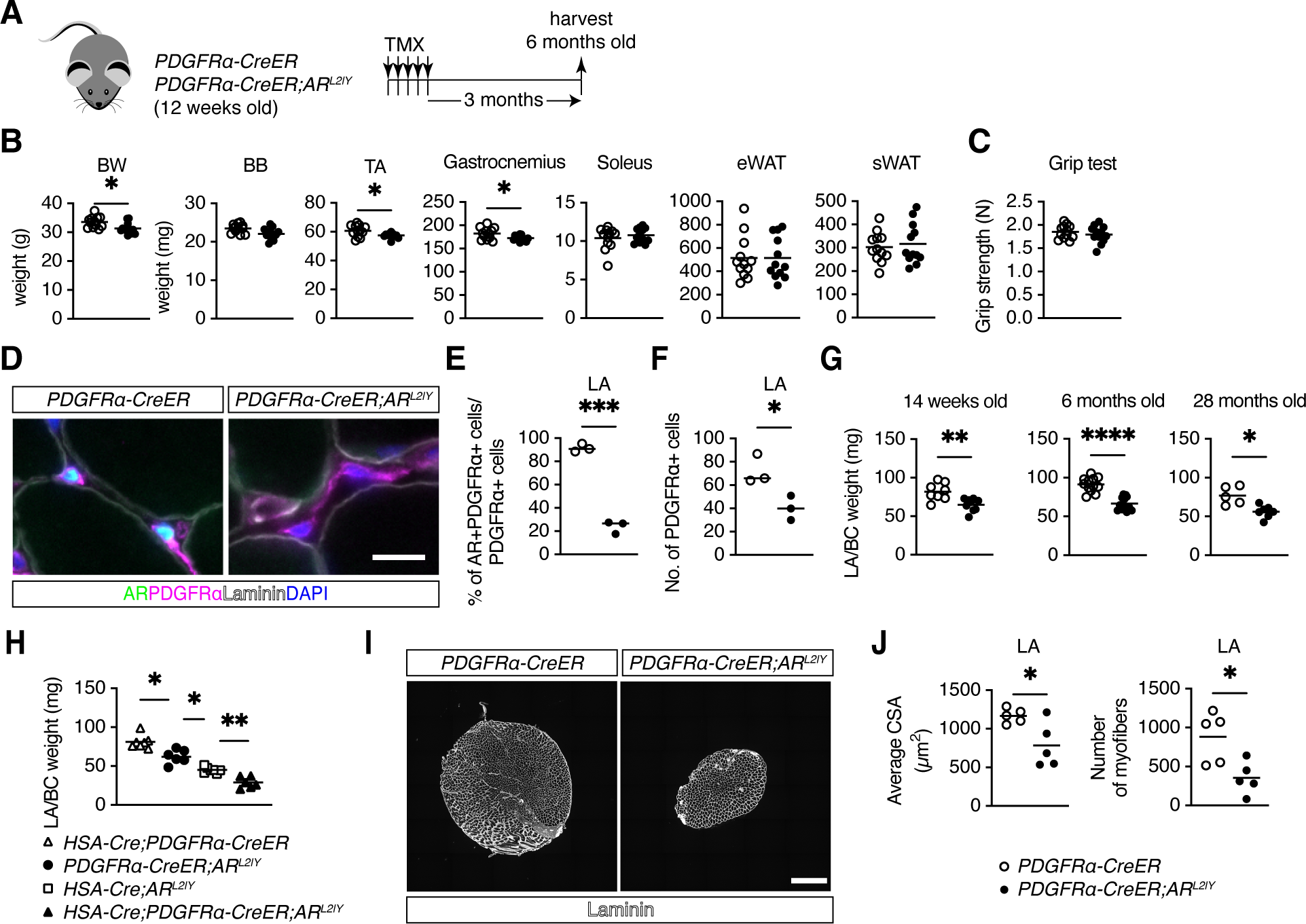
Atrophy of skeletal muscles in mesenchymal progenitor-specific AR-deficient mice. (A) Experimental design for tamoxifen (TMX) treatment followed by sample harvest in mature adult (6 months old) mice. (B and C) Quantification of body weight (BW), muscle mass, epididymal white adipose tissue (eWAT) mass, and subcutaneous WAT (sWAT) mass (B), and grip strength (C) in 6-month-old mice (n = 12). TA, tibialis anterior; BB, biceps brachii. Unpaired t-test with Welch’s correction. (D) Immunofluorescence staining of AR, PDGFRα, and laminin with DAPI in levator ani (LA) muscles of control (left) and mutant (right) mice. Scale bar, 10 µm. (E) Quantification of AR+ cells among PDGFRα+ cells in LA muscles of control and mutant mice (n = 3). The two-sample test for equality of proportions. (F) Quantification of PDGFRα+ cells in LA muscles of control and mutant mice (n = 3). Unpaired t-test with Welch’s correction. (G) Mass of LA and bulbocavernosus (BC) muscles in 14-week-(n = 8 control; n = 9 mutant mice), 6-month-(n = 12), and 28-month-old mice (n = 5 control; n = 7 mutant mice). Unpaired t-test with Welch’s correction. (H) Mass of LA/BC muscles in single (*HSA-Cre* or *PDGFRα-CreER*) and double Cre mice (n = 7 *HSA-Cre;PDGFRα-CreER* and *HSA-Cre;AR^L2/Y^*; n = 6 mutant and *HSA-Cre;PDGFRα-CreER;AR^L2/Y^* mice). Brown-Forsythe and Welch’s ANOVA test with Dunnett’s T3 multiple comparison test. (I) Immunofluorescence staining of laminin in LA muscles of control (left) and mutant mice (right). Scale bar, 500 µm. (J) Average cross-sectional area (CSA) (left) and the number of myofibers (right) in control and mutant mice (n = 5). Unpaired t-test with Welch’s correction: *p < 0.05, **p < 0.01, ***p < 0.001, ****p < 0.0001.

Additionally, the weights of the entire body, hindlimb muscles, and white adipose tissues were compared between aged (28 months old) control and mutant mice (Fig S2J). The aged mutant mice did not exhibit any significant differences from the aged control mice (Fig. S2K and S2L). Therefore, our findings demonstrate that AR deficiency in mesenchymal progenitors affects skeletal muscle mass in mature adult mice only, not young or aged mice, primarily in the limbs.

### Mesenchymal progenitor-specific AR ablation induced atrophy of LA/BC muscles

The perineal LA/BC skeletal muscle complex exhibits increased responsiveness to androgens compared with other striated muscles (26). Because this perineal muscle complex also has higher AR expression (14), we examined the impact of AR ablation on mesenchymal progenitors of LA/BC skeletal muscles. We confirmed efficient ablation of AR in PDGFRα+ cells in LA muscles as well as limb muscles (Fig. 2D, E). We also found that AR depletion in mesenchymal progenitors in LA muscles led to a decrease in the number of PDGFRα+ cells (Fig. 2F). At 14 weeks, 6 months, and 28 months of age, the LA/BC mass was consistently lower in the mutant mice than control mice (Fig. 2G). Because AR ablation in myofibers using a promoter region derived from the human α-skeletal actin (HSA) gene (*HSA-Cre;AR^L2/Y^* mice) also leads to reduced LA/BC mass (27), we generated *HSA-Cre;PDGFRα-CreER;AR^L2/Y^*mice to examine whether the weight reduction depended on AR in myofibers. Interestingly, the LA/BC weight was lower in *HSA-Cre;AR^L2/Y^* mice than in *PDGFRα-CreER;AR^L2/Y^* mice (Fig. 2H). Moreover, *HSA-Cre;PDGFRα-CreER;AR^L2/Y^*mice had the lowest LA/BC mass compared with *HSA-Cre;AR^L2/Y^* and *PDGFRα-CreER;AR^L2/Y^* mice (Fig. 2G). These findings suggest that AR expression in both myofibers and mesenchymal progenitors independently regulates the mass of perineal muscles. Examination of the cross-sectional area (CSA) of LA muscles by laminin staining (Fig. 2I) revealed a lower myofiber size and fewer myofibers in the mutant mice than control mice (Fig. 2J). In summary, AR expression in mesenchymal progenitors independently regulates the mass of perineal muscles, separate from its influence on myofibers.

### AR regulated genes related to extracellular matrix organization in mesenchymal progenitors

To determine the effects of AR on gene expression, we performed RNA-seq in PDGFRα+ cells from the LA/BC muscles of both control and mutant mice (Fig. 3A). Using a previously established gating strategy for isolating mesenchymal progenitors (CD45− CD31− Vcam1− PDGFRα+) (12) (Fig. S3A), we isolated 300 PDGFRα+ cells from LA/BC muscles for RNA-seq. We detected more than 100,000 genes in these cells, indicating sufficient read depth for further analysis (Fig. S3B). The principal component analysis (PCA) plot showed separation between the control and mutant samples (Fig. S3C). We identified 80 upregulated and 76 downregulated genes, including AR, in PDGFRα+ cells from the mutant mice (Fig. S3D). The upregulated genes were enriched in the Gene Ontology (GO) terms including regulation of cell death and cell motility (Fig. 3B). On the other hand, the downregulated genes were enriched in the GO terms related to extracellular matrix (ECM) organization and regulation of proteolysis (Fig. 3C).

**Figure 3.**
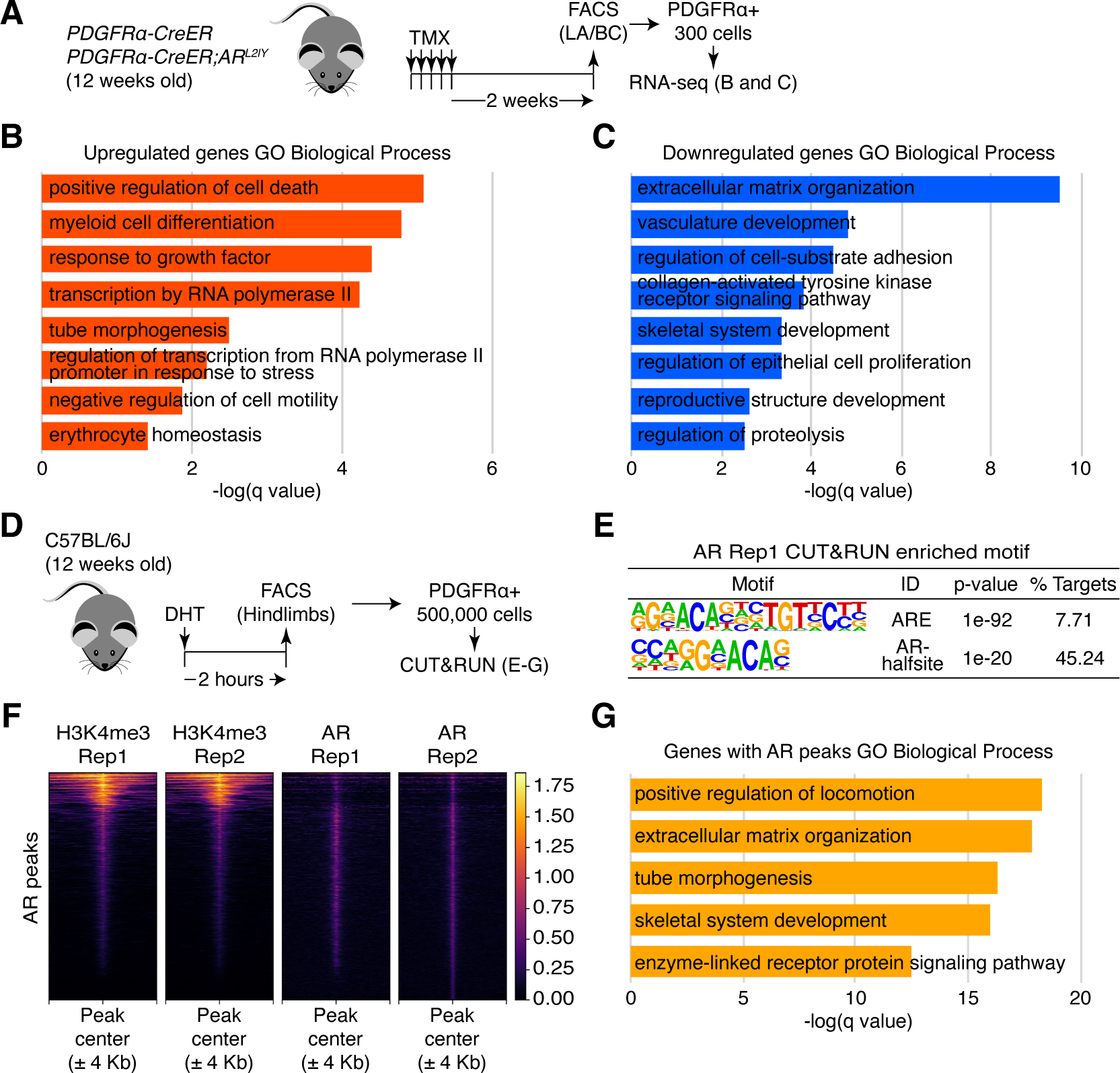
Functional and genomic targets of AR in mesenchymal progenitors of skeletal muscles. (A) Experimental design for tamoxifen (TMX) treatment followed by fluorescence-activated cell sorting (FACS) from levator ani and bulbocavernosus (LA/BC) muscles for RNA-seq. (B and C) Top Gene Ontology (GO) Biological Process terms associated with the upregulated (B) and downregulated (C) genes in PDGFRα+ cells from mutant mice. (D) Experimental design for dihydrotestosterone (DHT) treatment followed by FACS and CUT&RUN. (E) Enriched motifs within AR CUT&RUN peaks. (F) Heatmaps of mean H3K4me3 and AR CUT&RUN counts per million (CPM) normalized read coverage around AR CUT&RUN peaks for individual replicates. (G) Top GO Biological Process terms enriched within AR CUT&RUN peak-associated genes.

To determine the AR target genes, we used CUT&RUN method for AR and trimethylated lysine 4 from histone H3 (H3K4me3), which is enrich in active promoters, in mesenchymal progenitors isolated from C57BL/6J male mice (Fig 3D), at 2 hours after treatment with dihydrotestosterone (DHT), which was confirmed to induce AR expression in skeletal muscles (Fig. S3E, S3F). We detected total 6542 peaks of AR in mesenchymal progenitors in skeletal muscles. The AR peaks were primarily distributed in intron or intergenic regions (Fig. S3G), similar to previous reports with AR ChIP-seq in muscle and non-muscle tissues (28, 29). We also found the androgen response elements (ARE) in these peaks as the most enriched motif as well as AR half-site (Fig. 3E, S3H). Moreover, AR CUT&RUN was validated by peaks found in the *Fkbp5* locus (Fig. S3I), where canonical AR binding sites are located for several tissues (28). By profiling AR peaks with AR and H3K4me3 reads, we found that AR binding sites were present mainly not in promoter region (Fig. 3F). Furthermore, AR binding events were enriched for regulation of locomotion and ECM organization GO terms (Fig. 3G). These results indicate that AR in mesenchymal progenitors directly regulate the transcriptions including ECM organization genes.

### AR in mesenchymal progenitors regulated *Igf1* expression

To further investigate the gene regulation by AR in mesenchymal progenitors, we combined the RNA-seq and CUT&RUN analysis. We found 39 genes which were downregulated in AR-deficient mesenchymal progenitors and have AR peaks annotated by a nearest transcription start sites (TSS) analysis (Fig. 4A). Top enriched GO term of these genes was ECM organization (Fig. 4B), including Collagen family (*Col1a1*, *Col3a1*, *Col4a1*, *Col4a2*, *Col5a1*, *Col5a2*, and *Col6a2*, Fig. S4A and S4B), and *Igf1* (Fig. 4C), which is critical for maintaining muscle mass (30). We also found the bindings of AR with ARE and peaks of H3K4me3 in upstream of *Igf1* TSS (Fig. 4D). Moreover, AR peaks were found with ARE in *Ar* gene locus (Fig. S4C), indicating the regulation of *Ar* by AR in mesenchymal progenitors as other cells (31). To further investigate the effects of androgen and AR on *Igf1* expression, we isolated PDGFRα+ cells from C57BL/6J male mice treated with DHT (Fig. 4E). *Igf1* expression was upregulated in PDGFRα+ cells after DHT administration (Fig. 4F). We also found reduction of IGF1 protein in the mutant LA muscles (Fig. 4G and 4H). Finally, to determine whether the reduction in muscle mass in the mutant mice was due to decreased IGF1 protein, we injected recombinant IGF1 into the LA/BC muscles of the mutant mice just before TMX administration (Fig. 4I). We found that the mass of the LA/BC muscles in IGF1-treated mutant mice increased compared to saline-injected mutant mice, reaching levels comparable to those of saline-injected control mice (Fig. 4J).

**Figure 4.**
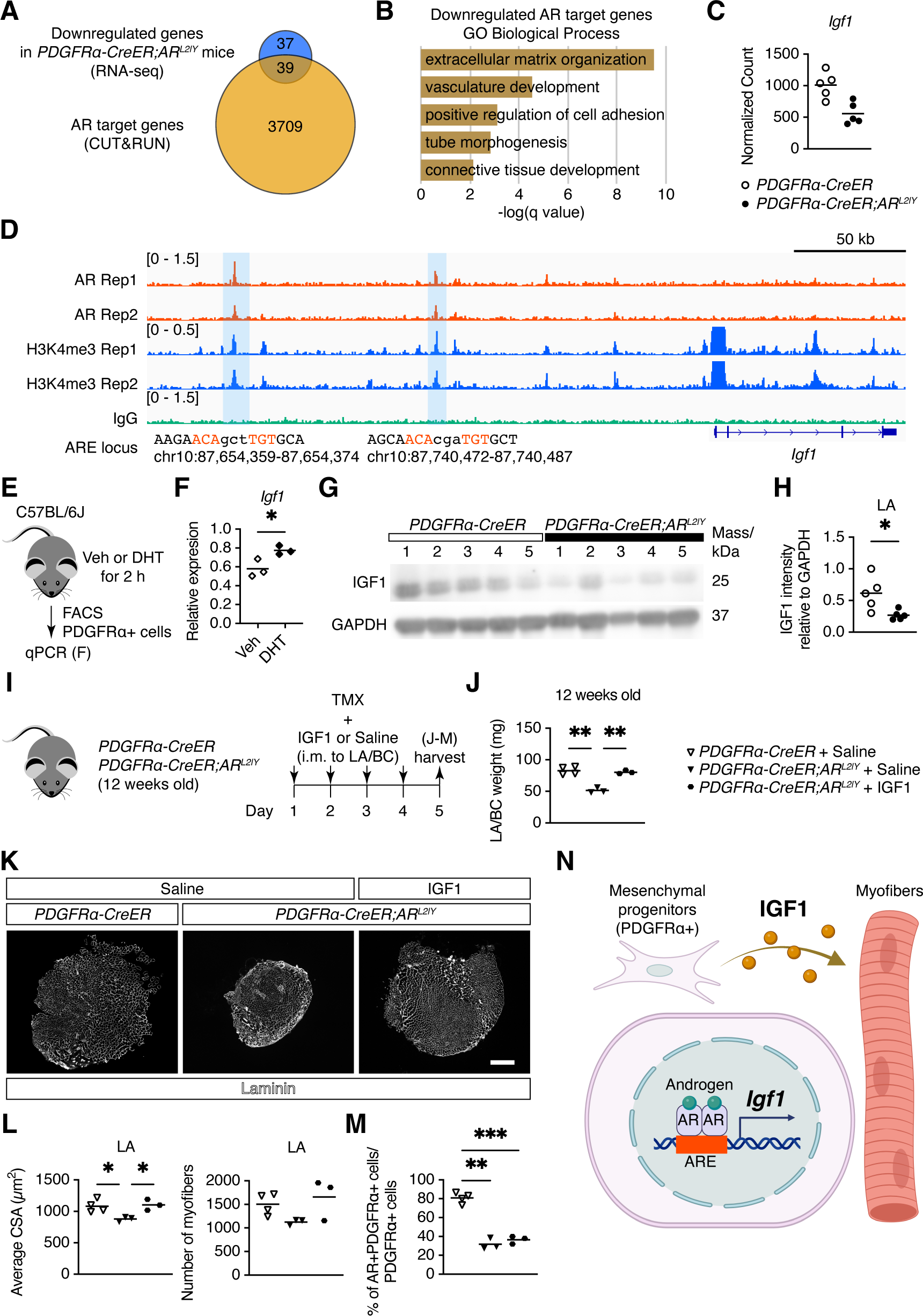
Regulation of *Igf1* in mesenchymal progenitors by AR. (A) Overlap of downregulated genes in PDGFRα+ cells from mutant mice and AR CUT&RUN peak-associated genes. (B) Top Gene Ontology (GO) Biological Process terms enriched within overlapped genes. (C) Normalized gene counts for *Igf1* obtained by RNA-seq. (D) AR and H3K4me3 CUT&RUN peaks with androgen response elements (ARE) at *Igf1* locus. The AREs are shown in red. Top left number is the y-axis range in counts per million (CPM). (E) Experimental design for qPCR after treatment with dihydrotestosterone (DHT). (F) qPCR analysis of *Igf1* in PDGFRα+ cells after DHT treatment (n = 3 mice/condition). Unpaired t-test with Welch’s correction. (G and H) Western blotting of IGF1 and GAPDH (G) and their quantification (H) in levator ani (LA) muscles from control and mutant mice (n = 5). Unpaired t-test with Welch’s correction. (I) Experimental design for intramuscular injection (i.m.) of IGF1 to LA and bulbocavernosus (BC) muscles, followed by tamoxifen (TMX) treatment and sample harvest in 12-week-old control and mutant mice. (J) Mass of LA/BC muscles in saline-or IGF1-injected control and mutant mice (n = 4 control, n = 3 mutant mice with saline or IGF1). Brown-Forsythe and Welch’s ANOVA test with Dunnett’s T3 multiple comparison test. (K) Immunofluorescence staining of laminin in LA muscles following saline injection in control mice (left), saline injection in mutant mice (middle), and IGF1 injection in mutant mice (right). Scale bar, 500 µm. (L) Average cross-sectional area (CSA) (left) and the number of myofibers (right) in saline-or IGF1-injected control and mutant mice (n = 4 control, n = 3 mutant mice with saline or IGF1). Brown-Forsythe and Welch’s ANOVA test with Dunnett’s T3 multiple comparison test. (M) Quantification of AR+ cells among PDGFRα+ cells in LA muscles of saline-or IGF1-injected control and mutant mice (n = 4 control, n = 3 mutant mice with saline or IGF1). The three-sample test for equality of proportions: *p < 0.05, **p < 0.01. ***p < 0.001. (N) Regulation of *Igf1* by AR in mesenchymal progenitors of skeletal muscle. Created with BioRender.com.

Immunohistochemical staining of laminin in LA muscles (Fig. 4K) revealed a significant reduction in muscle CSA of saline-injected mutant mice, which was prevented by IGF1 injection (Fig. 4L). Additionally, the number of myofibers tended to decline following TMX administration, and this trend was also mitigated by IGF1 injection (Fig. 4L). We also confirmed that AR was efficiently ablated as early as 4 days after TMX treatment, and that IGF1 injection did not impact the ablation efficiency (Fig. 4M). These findings indicate that IGF1 injection effectively prevents the atrophy of LA/BC muscles in mutant mice. Taken together, AR in mesenchymal progenitors regulates *Igf1* expression controlling skeletal muscle mass.

## Discussion

In the present study, we demonstrated that male mice with specific AR ablation in mesenchymal progenitors showed lower skeletal muscle mass in the limbs only at the mature adult stage, but not at a younger stage (14 weeks old). Taken together with previous report demonstrating that mice with AR ablation specifically in fast-twitch fibers lack muscle mass reduction at 13 weeks old but exhibit this reduction at 12–13 months old (23), there may be an effect that requires time to develop. Additionally, we demonstrated that the number of Pax7+ muscle stem cells was similar between adult control and mutant mice. This is in contrast to the observed reduction in myoblast proliferation following deletion of AR in the mesenchyme of BC muscles during development (16), providing evidence that AR in mesenchymal progenitors plays distinct roles during development, adulthood, and aging in mice. We also showed that specific ablation of AR in mesenchymal progenitors does not affect the fiber types of slow-twitch muscles. Depletion of mesenchymal progenitors results in a slight increase in the proportion of slow fibers (type I) predominantly in fast-twitch TA muscles (12), suggesting that factors other than AR in mesenchymal progenitors influence muscle fiber types. Muscle strength was either unaffected or only slightly affected in myofiber-specific AR ablation mice (32), suggesting that non-myogenic cells expressing AR may play a role in mediating muscle strength. However, grip strengths for forelimbs were not affected in the mutant mice in all experimental groups conducted in present study. Because the conventional grip strength test is influenced by various factors, including inconsistent procedures and the motivation of the mice (33), more sensitive methods for assessing muscle function, such as measuring in situ maximum isometric torque, and active and passive force in skinned fibers (34), could detect subtle differences in muscle strength in the mutant mice.

PDGFRα+ cells are the origin of ectopic adipocytes in skeletal muscles (9, 10). However, the specific molecular mechanisms underlying fatty infiltration of muscles remain largely unexplored (35). In general, androgen and AR signaling seem to have inhibitory effects on adipogenesis (36). Serum androgens are linked to reduced levels of fatty infiltration of skeletal muscles (37). In contrast, deprivation of androgens leads to an increase in fatty infiltration of skeletal muscle in men (38). Moreover, transgenic mice expressing AR induced by collagen type I alpha 1, which is also expressed in mesenchymal progenitors (39), have reduced body weight and fat mass (40). Additionally, HFD-induced fat mass deposition in skeletal muscles was increased by a low androgen level caused by orchidectomy (41). However, our results indicate that AR deficiency in mesenchymal progenitors does not affect adipogenesis in skeletal muscle even after HFD feeding or glycerol injection. An indirect androgen pathway may also regulate adipogenesis of mesenchymal progenitors in skeletal muscles. Indeed, recent research indicates that androgens can modulate macrophage polarization, subsequently affecting adipocyte differentiation (42). Further studies are needed to gain a comprehensive understanding of the mechanisms by which androgens and AR signaling influence fatty infiltration of muscle.

We observed a modest reduction in the muscle weight of the mutant mice at the mature adult age (24 weeks old), but this effect was not observed in aged mice (28 months old). Because selective ablation of AR in myofibers does not lead to reduced muscle mass even in 40-week-old mice (27), the role of AR in mesenchymal progenitors is more important than its role in myofibers in the context of muscle maintenance, at least at the mature adult age. The timing of AR ablation could impact skeletal muscle mass, given the variation in initiation times for AR ablation (tamoxifen treatment at 12 weeks old for analysis at 14 and 24 weeks, and at 8 weeks old for analysis at 28 months). Experiments involving AR ablation at different ages, such as TMX treatment at young (12 weeks old), mature adult (24 weeks old) and aged (24 months old) mice, would help to clarify whether these results are influenced by age or the duration of AR ablation. A decrease in testosterone concentration, even in 1-year-old mice compared with young 12-week-old mice (43), may contribute to muscle mass loss, potentially masking any subtle differences between the aged control and mutant mice. Due to the reduction in muscle mass observed in 2-year-old mice with AR depletion specifically in fast-twitch fibers (23), AR expression in muscle fibers may play a crucial role in regulating muscle metabolism and quality, which help maintain muscle mass in aged mice.

The perineal skeletal muscles LA and BC are highly androgen sensitive. Muscle fiber-specific AR knockout results in a more pronounced mass reduction in LA/BC muscles compared with limb muscles (26, 27), which is consistent with our present study demonstrating a severe reduction in LA/BC muscles in mice with AR ablation in mesenchymal progenitors. We also observed that the decrease in LA/BC weight in mice with AR ablation in mesenchymal progenitors was less pronounced than in those with AR ablation in myofibers. Furthermore, the double Cre mice (*HSA-Cre;PDGFRα-CreER;AR^L2/Y^*), with ablation of AR in both myofibers and mesenchymal progenitors, exhibited a greater reduction in LA/BC mass compared with single Cre mice (*HSA-Cre;AR^L2/Y^* or *PDGFRα-CreER;AR^L2/Y^*). Overexpression of AR in skeletal muscle fibers using HSA promoter (44, 45) does not rescue the atrophy of LA muscles of testicular feminization mutation (*Ar^Tfm^*) male rat (45), supporting a separate role of AR in mesenchymal progenitors for maintaining perineal muscle mass. Taken together, AR expression in mesenchymal progenitors independently regulates the mass of perineal muscles, separate from the influence of AR in myofibers. The distinct reactions observed in perineal and limb muscles could be attributed to a greater abundance of myonuclei expressing AR in perineal muscles compared with limb muscles (14). Nevertheless, our findings revealed that the majority of PDGFRα+ cells exhibited AR expression in both limb and perineal muscles, suggesting an additional mechanism by which AR in mesenchymal progenitors regulates muscle mass and other mesenchymal progenitor functions.

AR ablation in PDGFRα+ cells in adult mice led to a reduction in both muscle CSA and the number of myofibers in LA muscles. Additionally, our findings demonstrated that IGF1 injection into LA muscles protected against these reductions. Since age-related muscle loss, or sarcopenia, is primarily characterized by a decrease in both the number and area of myofibers (46), it is possible that the age-related decline in androgen levels, which results in reduced AR expression in mesenchymal progenitors, contributes to these changes. Supporting this idea, muscle-specific IGF1 overexpression in aged mice preserves myofiber numbers (47), while IGF1 receptor knockout in myofibers leads to a reduction in myofiber count (48), suggesting a critical role for the AR/IGF1 axis in mesenchymal progenitors for maintaining myofibers. The reduction of myofibers during aging is driven by various mechanisms, including satellite cell depletion, altered protein synthesis, and increased muscle protein degradation (49). It’s important to note that the sensitivity of perineal muscles to androgens differs from that of limb muscles, suggesting that distinct AR-mediated mechanisms may be responsible for maintaining myofibers in these different muscle groups. Although our study found that genes related to cell death, which can lead to decreased protein synthesis, were upregulated in mesenchymal progenitors of the perineal LA/BC muscles following AR ablation, additional research is needed to elucidate the molecular mechanisms by which AR/IGF1 axis dysfunction in mesenchymal progenitors contributes to myofiber loss.

We also found that several ECM genes, including members of the collagen family, were downregulated in AR-deficient mesenchymal progenitors, with AR peaks identified by a nearest TSS analysis. These ECM components are primarily produced by PDGFRα+ cells in both healthy and fibrotic skeletal muscles (39). AR expression promotes fibrosis in the heart, kidney, and ovary (50–52), indicating that androgen/AR signaling governs ECM remodeling. Moreover, downregulation of AR in PDGFRα+ fibroblasts from squamous cell carcinomas induces the activation of cancer-associated fibroblasts (CAF), which remodel the ECM within the tumor microenvironment (53). This suggests a general role for AR in PDGFRα+ cells under both physiological and pathological conditions. Given that muscle diseases involving ECM remodeling, such as muscular dystrophy and sarcopenia, are often accompanied by muscle atrophy (54), it is plausible that ECM reduction due to AR deficiency contributes to skeletal muscle atrophy. Further studies are needed to elucidate the relationship between AR, ECM, and skeletal muscle plasticity.

The signaling pathway involving IGF1 plays a crucial role in both the developmental growth and steady-state maintenance of skeletal muscle mass (55, 56). IGF1 is produced primarily in the liver (57) but local production, including in skeletal muscles, is induced by androgen treatment. The administration of testosterone to older men led to an increase in IGF1 protein expression in the limb (vastus lateralis) muscles (58). *Igf1* expression is decreased in the LA/BC muscles of mice with skeletal muscle-specific AR ablation (27). Dubois et al. has reported indirect androgen regulation of IGF1 via non myocytic AR because Myod-Cre-driven AR-deficient mice showed no reduction in the *Igf1* transcript level, and *Igf1* expression was decreased after castration in skeletal muscles (17). We demonstrated that AR in mesenchymal progenitors in skeletal muscles regulated *Igf1* expression, governing muscle mass. Moreover, activation of Akt signaling via the expression of Bmp3b in mesenchymal progenitors (12) suggests a connection with Igf1, highlighting the complex interplay of signaling pathways in the maintenance of skeletal muscle mass.

A limitation of our study in that we focused on male mice for studying AR in mesenchymal progenitors of skeletal muscles. Because serum levels of androgens, including testosterone and DHT, are also measurable in female mice (43), androgen/AR signaling could potentially affect female skeletal muscles as well. Furthermore, *AR* was not differentially expressed between males and females of human skeletal muscles (59), indicating potential roles of AR in female skeletal muscles. On the other hand, genomic AR knockout mice show impaired skeletal muscle development and function in males, but not females (5). Although muscle fiber-specific AR knockout in female mice limitedly impacted on the expression of transcripts (6) and muscle mass (23), more comparative work between male and female mice will be necessary to determine whether AR plays important roles in female mesenchymal progenitors, as the function of AR in these cells may differ from its role in female skeletal muscle fibers.

In summary, our results show that AR in mesenchymal progenitors regulates skeletal muscle mass via *Igf1* expression (Fig. 4N). Further investigations into the limb muscles of aged mice and humans are required to confirm whether combined treatment with androgens and IGF1 could be effective in treating muscle atrophy conditions such as sarcopenia.

## Materials and Methods

### Animals

*PDGFRα-CreER* mice (19), *HSA-Cre* mice (60), and C57BL/6J male mice were purchased from Jackson Laboratories (Strain #: 018280, 006149, and 000664, respectively). *AR^L2/Y^* mice have been described (20). *PDGFRα-CreER* male mice were crossed with *AR^L2/+^* female mice to generate *PDGFRα-CreER;AR^L2/Y^* male mutant mice. Male littermates with the *PDGFRα-CreER;AR^+/Y^* genotype were used as control mice. To generate *HSA-Cre;PDGFRα-CreER;AR^L2/Y^* male mice, *HSA-Cre* male mice were crossed with *PDGFRα-CreER;AR^L2/+^* female mice. All mice were housed in a specific pathogen-free facility under climate-controlled conditions and a 12-hour light/dark cycle and were provided water and a standard diet *ad libitum*. In both control and mutant mice, 100 µL of 20 mg/mL tamoxifen (TMX, Sigma-Aldrich, Cat# T5648), dissolved in corn oil (Sigma-Aldrich, Cat# C8267), was injected intraperitoneally for 5 consecutive days at 8 or 12 weeks of age. A high-fat diet (HFD, CLEA Japan, Cat# HFD32) was given *ad libitum* for 8 weeks to induce obesity in mice. The diet was refreshed every 2 days. To induce fatty infiltration of skeletal muscle, 100 µL of 50% v/v glycerol was injected into tibialis anterior (TA) muscles under anesthesia using isoflurane (61). To induce AR signaling, vehicle (Veh) or dihydrotestosterone (DHT or stanolone, 2.5 mg/mouse; Tokyo Chemical Industry, Cat# A0462), dissolved in absolute ethanol and diluted in 0.3% w/v hydroxypropyl cellulose (Wako, Cat# 085-07932) in PBS, was injected subcutaneously into C57BL/6J male mice at 2 hours before sampling. In *PDGFRα-CreER;AR^L2/Y^*male mutant mice, 10 µg/50 µL of recombinant IGF1 (PeproTech, Cat# AF-100-11) dissolved in saline (Otsuka, Cat# 3311401A2026) or 50 µL of saline was injected directly into LA/BC muscles for 4 consecutive days to keep the concentration of IGF1 in the muscles at 12 weeks of age (62). The same amount of saline was injected into LA/BC muscles of male littermates with the *PDGFRα-CreER;AR^+/Y^*genotype for the control. TMX (100 µL of 20 mg/mL) was injected intraperitoneally, just after injection of IGF1 or saline, for 4 consecutive days. All animal experiments were approved by the Animal Experiment Committee of Ehime University (Approval No. 37A1-1·16) and were conducted in full compliance with the Guidelines for Animal Experiments at Ehime University.

### Grip strength

The grip strength of the forelimb was measured using a strain gauge (Melquest, Cat# GPM-100B). The maximal force was determined after 10 measurements. The mean value recorded by two independent experimenters was reported as the grip strength of each mouse.

### Immunofluorescence staining and microscopy

For immunofluorescence staining, muscles were snap frozen in liquid nitrogen-chilled isopentane. Cryosections of 10 µm thickness were prepared for immunofluorescence staining. Sections were fixed in 4% paraformaldehyde in PBS for 5 min, blocked in 5% goat serum (Gibco, Cat# 16210-064) in PBS for 60 min, and incubated with primary antibodies overnight at 4°C. For staining of myosin heavy chain (Myh), sections were air-dried and blocked with M.O.M. reagent (Vector Laboratories, Cat# MKB-2213-1) at room temperature for 60 min.

Primary antibodies for Myh diluted in 1% BSA/PBS were incubated at 37°C for 45 min. The antibodies used for immunofluorescence staining were anti-AR (Abcam, Cat# ab108341, RRID:AB_10865716; 1/100), anti-PDGFRA (R&D Systems, Cat# AF1062, RRID:AB_2236897; 1/80), anti-Laminin α2 (Santa Cruz Biotechnology, Cat# sc-59854, RRID:AB_784266; 1/400), anti-Pax7 (DSHB, Cat# PAX7, RRID:AB_528428; 1/5), anti-Laminin (Sigma-Aldrich, Cat# L9393, RRID:AB_477163; 1/500), anti-Myh7 (DSHB, Cat# BA-D5, RRID:AB_2235587, supernatant; 1/100), anti-Myh2 (DSHB, Cat# SC-71, RRID:AB_2147165, supernatant; 1/100), anti-Perilipin (Sigma-Aldrich, Cat# P1873, RRID:AB_532267; 1/250), Alexa fluor 488 goat anti-rabbit IgG (Thermo Fisher Scientific, Cat# A-11008; RRID:AB_143165; 1/1000), Alexa fluor 555 donkey anti-goat IgG (Thermo Fisher Scientific, Cat# A-21432, RRID:AB_2535853; 1/1000), Alexa fluor 647 goat anti-rat IgG (Thermo Fisher Scientific, Cat# A-21247, RRID:AB_141778; 1/1000), Alexa fluor 568 goat anti-mouse IgG1 (Thermo Fisher Scientific, Cat# A-21124, RRID: AB_2535766; 1/1000), Alexa fluor 488 goat anti-mouse IgG2b (Thermo Fisher Scientific, Cat# A-21141, RRID:AB_2535778; 1/1000), Alexa fluor 568 goat anti-rabbit IgG (Thermo Fisher Scientific, Cat# A-11011, RRID:AB_143157; 1/1000), and Alexa fluor 647 goat anti-rabbit IgG (Thermo Fisher Scientific, Cat# A27040; RRID:AB_143165; 1/1000). Nuclei were counterstained with DAPI. Entire cross sections of fluorescence images were photographed using BIOREVO (Keyence) or Axio Observer7 (Zeiss).

### Quantification of muscle sections

To quantify the number, cross-sectional area (CSA), and minimum Ferret diameter of the myofibers, captured images of entire cross sections were analyzed using Fiji (v2.3.0) (https://imagej.net/Fiji). The myofiber area was identified by the laminin+ signals. The region of interest (ROI) was set within the laminin+ areas. The frequency of each muscle fiber type in the ROI was recorded by Trainable Weka Segmentation in Fiji after manual training. Small (< 150 µm^2^) or large (> 10,000 µm^2^) fibers were excluded for analyzing. The number of AR+PDGFRα+ mesenchymal progenitors, Pax7+ satellite cells, and the perilipin+ area was manually counted or measured using Fiji.

### Isolation of mesenchymal progenitors from skeletal muscle

Isolation of mononuclear cells from muscle was performed as described previously (63) with minor modifications. Briefly, muscles were chopped in PBS on ice and digested using 800 U/mL collagenase type 2 (Worthington, Cat# LS004177) in F10 medium (Gibco, Cat# 11550-043) supplemented with 10% horse serum (Gibco, Cat# 26050-088) and antibiotic–antimycotic (Gibco, Cat# 15240-062) for 60 min. Another digestion was conducted using 1000 U/mL collagenase type 2 and 11 U/mL dispase (Gibco, Cat# 17105-041) for 30 min. The supernatants were transferred to a 40 µm cell strainer (Falcon, Cat# 352340) to filter the debris and then centrifuged to yield a cell suspension. The following antibodies were used: FITC anti-CD31 (BioLegend, Cat# 102406; RRID:AB_312901; 1/100), FITC anti-CD45 (BioLegend, Cat# 103108, RRID:AB_312973; 1/100), PE anti-VCAM1 (Thermo Fisher Scientific, Cat# 12-1061-82, RRID:AB_2572573; 1/100), and APC anti-PDGFRα (R&D Systems, Cat# FAB1062A, RRID:AB_3073817; 1/30). The BD FACSAria II (RRID:SCR_018091) or BD FACSAria Fusion (BD Biosciences) was used for fluorescence-activated cell sorting (FACS), and the plot was analyzed by FlowJo v10.1r7 (BD Life Sciences, RRID:SCR_008520).

### RNA-seq and data analysis

For isolation of RNA from mesenchymal progenitors in LA/BC muscles, 300 PDGFRα+ cells were sorted into the wells of a 96-well plate containing 10 µL lysis buffer at 4°C. The lysis buffer was prepared by mixing 1 µL RNase inhibitor (Takara, Cat# 2313A) and 19 µL 10X lysis buffer (Takara, Cat# 635013) diluted to 1X with nuclease-free water. 3’ UTR RNA-seq was conducted using the CEL-Seq2 protocol (64), with the exception of employing the Second Strand Synthesis Module (New England Biolabs, Cat# E6111L) for double-stranded cDNA synthesis. The library was then amplified via 10 cycles of PCR without any sample pooling. Subsequently, the library was sequenced on the Illumina NovaSeq 6000, and quantitative analysis was performed using 81 bp of insert reads (Read2).

Cell barcode and UMI information from Read1 were extracted using UMI-tools (v1.1.2) (65) with the following command: “umi_tools extract -I read1.fastq --read2-in=read2.fastq --bc-pattern=NNNNNNNNNNCCCCCCCCC”. Adaptor and low-quality sequences were removed, and reads with a length less than 20 bp were discarded using TrimGalore (v0.6.7; http://www.bioinformatics.babraham.ac.uk/projects/trim_galore/). The processed reads were then mapped to the GRCm38 reference genome using HISAT2 (v2.2.1) (66). Read counts for each gene were obtained using featureCounts (v2.0.1) (67), and UMI duplications were removed using UMI-tools. Four cell barcodes, “ATGCAATGC,” “GACACGACA,” “CGATGCGAT,” and “TCTGGTCTG,” were assigned to each sample. Differentially expressed genes were identified using DESeq2 (v1.36.0) (68) with thresholds of |log2FC| > 1 and padj < 0.1. Metascape (69) was used for Gene Ontology (GO) enrichment analysis.

### CUT&RUN and data analysis

C57BL/6J male mice (10-12 weeks old) were treated with DHT at 2 hours before sampling for FACS. For isolation of chromatin from mesenchymal progenitors in hindlimbs muscles, 500,000 PDGFRα+ cells were sorted to 20%FBS, DMEM/F12 (Gibco, Cat# 10565-018), Ultroser G (Pall, Cat# 15950-017) supplemented with DHT (100 nM) to activate AR signaling during sorting. After sorting, the nuclei were prepared with ChIC/CUT&RUN Assay Kit (Active Motif, Cat# 53180) following the manufacturer’s protocol. For AR (n = 4), H3K4me3 (n = 2), and IgG control (n = 2) CUT&RUN, the following antibodies (1 µg/reaction) were used: anti-AR (Abcam, Cat# ab108341, RRID:AB_10865716), anti-H3K4me3 (Active Motif, Cat# 39916, RRID:AB_2687512), and rabbit negative control IgG (Active Motif, part of Cat# 53180). To obtain enriched DNA after the CUT&RUN reaction, 20 µL of DNA purification elution buffer was used instead of 55. The library was prepared using NEBNext Ultra II for DNA Library Preparation for Illumina (New England Biolabs, Cat# E7645L) with some modifications according to the Active Motif’s instruction. All reactions were performed by half volume. Constructed libraries were analyzed by 2100 Bioanalyzer using High Sensitivity DNA kit (Agilent Technologies, Cat# 5067-4626), and quantified by qPCR using KAPA Library Quantification Kits (Kapa Biosystems, Cat# KK4824). The library was sequenced on the Illumina NovaSeq 6000 for 51 cycles of paired-end sequencing.

Paired-end reads were trimmed with TrimGalore and mapped to mm10 using Bowtie2 (v2.4.5) (70) with the following options: --no-mixed --no-discordant. Multi or no mate reads were removed with samtools (v1.10) (71) view (-q 4 -F 0×2). To identify AR peaks across samples, BAM files were merged across biological replicates (n = 2) to call peaks with MACS2 (v2.2.7.1) (-q 0.05) (72). H3K4me3 peaks were called on individual BAM files using MACS2 (-q 0.0001). Merged BAM files for AR and IgG were used for downstream analysis (2 merged BAM files for AR and 1 merged BAM file for IgG). Motif enrichment analysis of AR peaks was performed with HOMER (v4.11) (73) findMotifs with -size 200 -mask options and AR peaks were annotated to mm10 genes using annotatePeaks (-size 200 -mask) from MACS2 summit files. Peaks distribution was annotated on mm10 (TxDb.Mmusculus.UCSC.mm10.knownGene, R package version 3.4.7.) using ChIPpeakAnno package (v3.30.1) (74, 75) with MACS2 narrowPeak files. Promoter region was defined as 2000 bp upstream to 500 bp downstream of transcription start sites (TSS). Common AR peaks-associated genes between replicates were identified with Venn diagrams tools (http://bioinformatics.psb.ugent.be/webtools/Venn/). GO enrichment analysis of common AR peaks-associated genes for GO Biological Process was performed with Metascape. To generate bigwig files, merged (AR or IgG) or individual (H3K4me3) BAM files were converted to CPM-normalized bigwig files using deepTools (v3.5.1) (76) bamCoverage (--binSize 100 --normalizeUsing CPM). Peaks were visualized using Integrative Genomics Viewer (IGV, v2.12.3) (77) with bigwig files. A count matrix for aggregation plot was generated by deepTools computeMatrix reference-point (--skipZeros) with AR and H3K4me3 bigwig files. Heatmaps were generated with count matrix files using deepTools plotHeatmap.

### RNA isolation and quantitative real-time PCR

C57BL/6J male mice (12 weeks old) were treated with Veh or DHT at 2 hours before sampling for FACS, as described above. Total RNA was extracted from freshly sorted 30,000 PDGFRα+ cells from LA/BC muscles using the RNeasy Plus Micro Kit (Qiagen, Cat# 74034) following the manufacturer’s protocol. Reverse transcription was performed using PrimeScript (Takara, Cat# RR036A) to synthesize cDNA from total RNA. Quantitative PCR (qPCR) was conducted in technical duplicate samples utilizing TB Green Premix Ex Taq II (Takara, Cat# RR820S) and the Thermal Cycler Dice (Takara, Cat# TP850). The expression levels of the target genes were normalized to that of *Rpl13a*. The primer sequences are listed in Table S1.

### Western blotting

LA muscles were dissolved in RIPA buffer (Fujifilm, Cat# 182-02451) supplemented with protease inhibitors (Nacalai Tesque, Cat# 25955-11). Tissue-extracts were separated by SDS-PAGE and transferred to polyvinylidene fluoride membranes. The membranes were blocked with 5% skim milk in TBS with Tween-20 (PBST), followed by incubation with anti-IGF1 (R&D Systems, Cat# AF791, RRID:AB_2248752; 1/2000) or anti-GAPDH antibodies (Cell Signaling Technology, Cat# 5174, RRID: AB_10622025; 1/1000) overnight at 4°C. The membranes were incubated with an anti-goat (Agilent, Cat# P0160, 1:5000) or anti-rabbit immunoglobulin/HRP secondary antibody (Agilent, Cat# P0448, RRID: AB_2617138; 1:5000) for 60 min at room temperature. Signals were detected using ECL Prime Western Blotting Detection Reagents (Amersham, Cat# RPN2232) on the Image Quant LAS 4000 system (GE Healthcare). Band images were quantified by Fiji.

### Statistical analysis

Prism 9 (GraphPad Software) or R (v4.0.5) (https://www.r-project.org) was used for the statistical analysis. Welch’s t-test was used to compare two groups. The two-or three-sample test for equality of proportions was performed using the test of equal or given proportions. To compare variables among three or more groups, Brown-Forsythe and Welch’s ANOVA test with Dunnett’s T3 multiple comparison test was used.

## Acknowledgments and funding sources

We are grateful to A. Nishio and Y. Sato for their technical support. We thank Y. Tanaka for the flow cytometry support, N. Tokunaga for the library preparations, and other members of the Division of Medical Research Support, the Advanced Research Support Center, Ehime University. This study was partly performed in the Cooperative Research Project Program in the Medical Institute of Bioregulation (MIB), Kyushu University. We also thank the Medical Research Center for High Depth Omics, MIB, Kyushu University for the sequencing, and RIST at Kyushu University for the high-performance computing. This work was supported in part by MEXT/JSPS KAKENHI (JP21K17568 to H.S.; JP22H03203 to Y.I.) and the HIRAKU-Global Program, which is funded by MEXT’s “Strategic Professional Development Program for Young Researchers” and Takeda Science Foundation (to H.S.).

## Data Availability

RNA-seq and CUT&RUN data have been deposited and are publicly available in the Gene Expression Omnibus as accession number GSE247378.

