## Supplemental information for "The androgen receptor in mesenchymal progenitors regulates skeletal muscle mass via *Igf1* expression in male mice"

**This PDF file includes:**

Figures S1 to S4
Tables S1

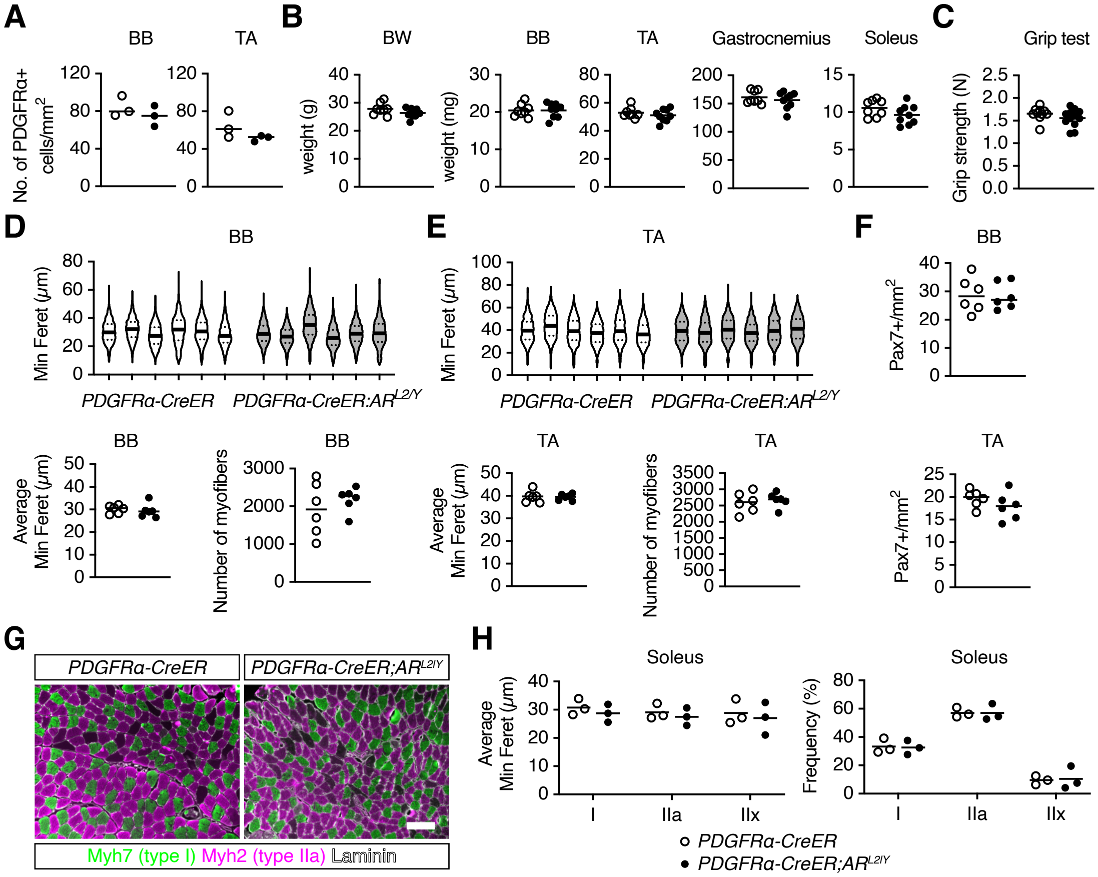


**Fig. S1.** Limited impact of AR deficiency in mesenchymal progenitors on intact limb skeletal muscles. (A) Quantification of PDGFRα+ cells in the biceps brachii (BB, left) and tibialis anterior (TA, right) muscles of control and mutant mice (n = 3). (B and C) Quantification of body weight (BW), muscle mass (B) and grip strength (C) at 2 weeks after tamoxifen treatment (n = 8 control; n = 9 mutant mice). (D and E) Violin plots of the minimum Ferret diameters of each individual (top) with their mean (bottom left) and quantification of the number of myofibers (bottom right) in the BB (D) and TA (E) muscles of control and mutant mice (n = 6). (F) Quantification of the number of Pax7+ cells in BB (top) and TA (bottom) muscles of control and mutant mice (n = 6). (G) Immunofluorescence staining of Myh7, Myh2, and laminin in the soleus muscles of control (left) and mutant mice (right). Scale bar, 100 µm. (H) Average minimum Ferret diameter (left) and frequency (right) of each fiber type in control and mutant mice (n = 3).

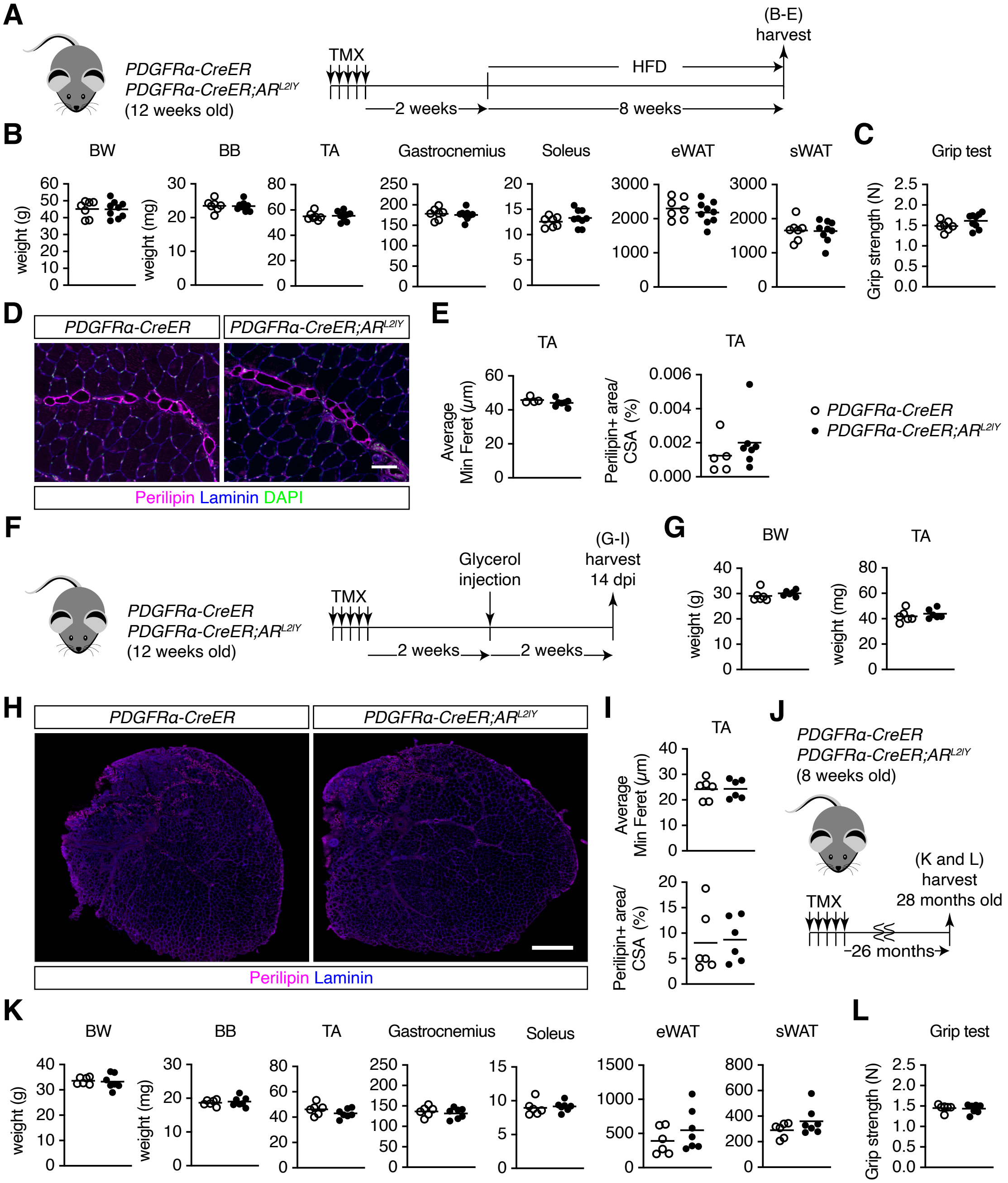


**Fig. S2.** Limited impact of AR deficiency in mesenchymal progenitors on adipogenesis in muscles. (A) Experimental design for tamoxifen (TMX) treatment followed by high fat diet (HFD) feeding. (B and C) Quantification of body weight (BW), muscle mass, epididymal white adipose tissue (eWAT) mass, and subcutaneous WAT (sWAT) mass (B), and grip strength (C) at 8 weeks after HFD feeding (n = 7 control; n = 9 mutant mice). TA, tibialis anterior; BB, biceps brachii. (D) Immunofluorescence staining of perilipin and laminin with DAPI in TA muscles of control (left) and mutant mice (right) after HFD feeding. Scale bar, 100 µm. (E) Average minimum Ferret diameter (left) and perilipin+ area (right) in TA muscles (n = 5 control; n = 7 mutant mice). (F) Experimental design for TMX treatment followed by glycerol injection. (G) Quantification of BW and TA muscle mass at 2 weeks after glycerol injection (n = 6). (H) Perilipin and laminin immunofluorescence staining in the TA muscles of control (left) and mutant mice (right) after glycerol injection. Scale bar, 500 µm. (I) Average minimum Ferret diameter (top) and perilipin+ area (bottom) of TA muscles after glycerol injection (n = 6). (J) Experimental design for TMX treatment followed by sample harvest in aged (28 months old) mice. (K and L) Quantification of BW, muscle mass, eWAT mass, sWAT mass (K), and grip strength (L) in 28-month-old mice (n = 6 control; n = 7 mutant mice).

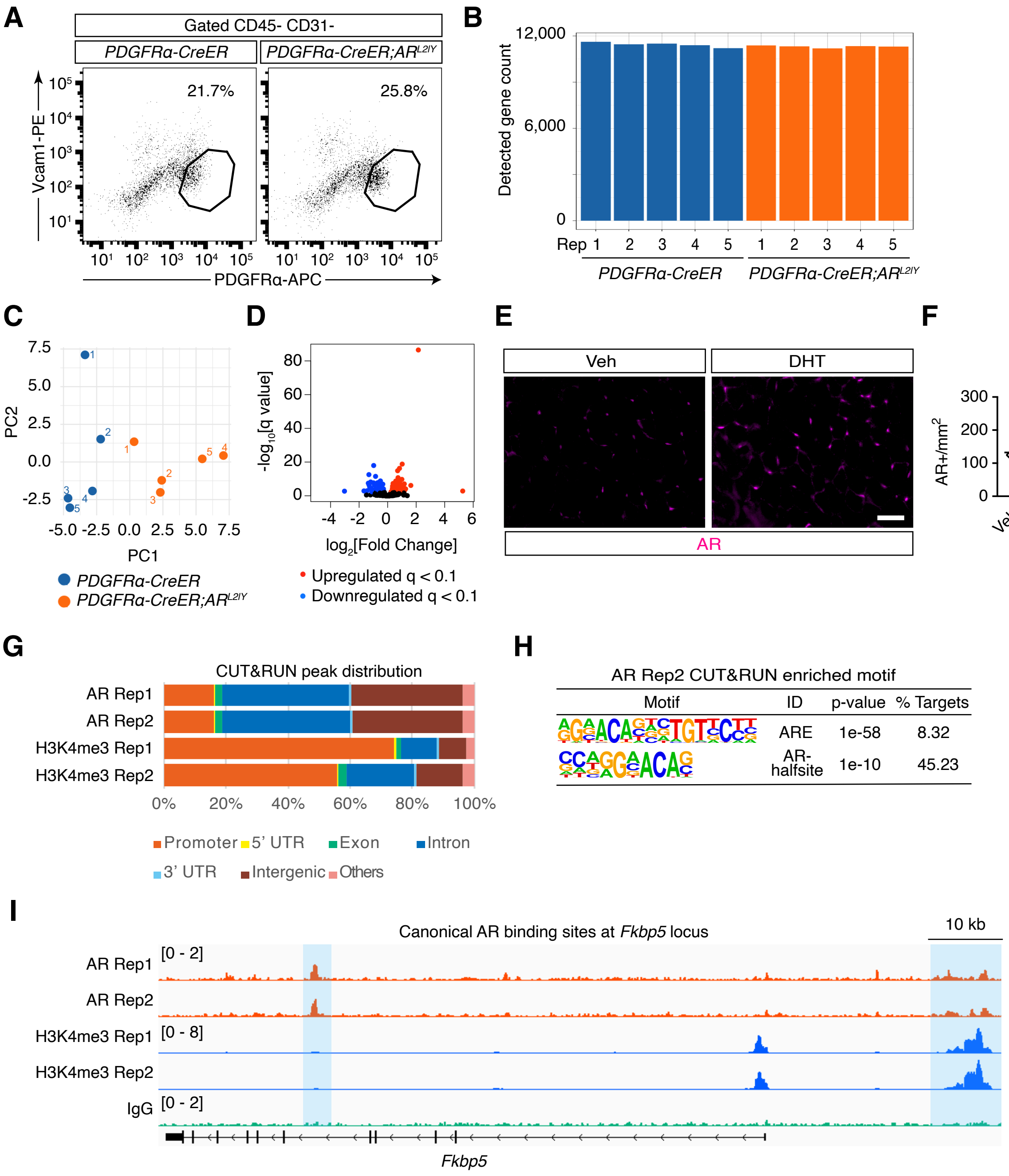


**Fig. S3.** Validation of RNA-seq and CUT&RUN in mesenchymal progenitors from skeletal muscles. (A) Gating strategy of fluorescence-activated cell sorting (FACS) for isolating mesenchymal progenitors (PDGFRα+ Vcam1− CD45− CD31−) for RNA-seq. (B) Gene counts detected in each RNA-seq sample (n = 5). (C) Principal component analysis (PCA) plot of the RNA-seq samples. (D) Volcano plot showing the $\log_2$ fold change (LFC, x-axis) and $-\log_{10}$ q-value (y-axis) of each gene in the differential expression analysis; red dots: DESeq2, LFC > 0, q < 0.1; blue dots: DESeq2, LFC < 0, q < 0.1. Positive and negative FC values indicate upregulated and downregulated genes, respectively, in PDGFRα+ cells from mutant mice. (E) AR immunofluorescence staining in the biceps brachii (BB) muscles 2 hours after subcutaneous injection of vehicle (Veh, left) or dihydrotestosterone (DHT, right). (F) Quantification of the number of AR+ cells in BB muscles of Veh or DHT treated mice (n = 3). (G) Distribution of AR and H3Kme3 CUT&RUN peaks. (H) Enriched motifs within AR CUT&RUN peaks. (I) AR CUT&RUN peaks at canonical AR binding sites at *Fkbp5* locus. Top left number is the y-axis range in counts per million (CPM).

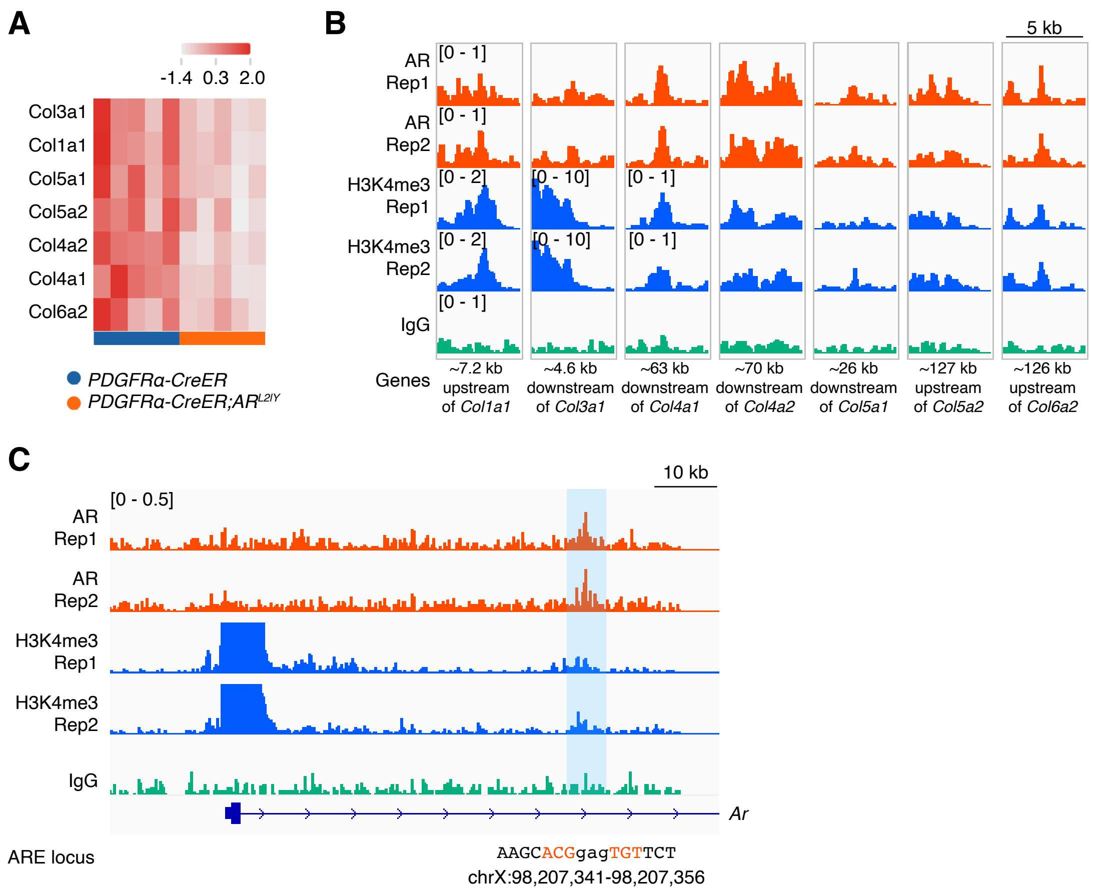


**Fig. S4.** Regulation of collagen family genes and *Ar* in mesenchymal progenitors by AR. (A) Heatmap showing the RNA-seq Z score of differentially expressed collagen family genes. (B) AR and H3K4me3 CUT&RUN peaks at collagen family genes. Top left number is the y-axis range in counts per million (CPM). (C) AR and H3K4me3 CUT&RUN peaks with androgen response elements (ARE) at *Ar* locus. The ARE is shown in red. Top left number is the y-axis range in CPM.

**Table S1.** Oligonucleotides used in this study.

| Oligo name | Sequence | Additional information |
|---|---|---|
| Igf1_F | TGGATGCTCTTCAGTTCGTG | qPCR |
| Igf1_R | CACTCATCCACAATGCCTGT | qPCR |
| Rpl13a_F | GTGGTCCCTGCTGCTCTCAAG | qPCR |
| Rpl13a_R | CGATAGTGCATCTTGGCCTTTT | qPCR |
